# Identification of dynamical changes of rabies transmission under quarantine: community-based measures towards rabies elimination

**DOI:** 10.1101/2023.05.17.541072

**Authors:** Kristyna Rysava, Michael J. Tildesley

## Abstract

Quarantine has been long used as a public health response to emerging infectious diseases, particularly at the onset of an epidemic when the infected proportion of a population remains identifiable and logistically tractable. In theory, the same logic should apply to low-incidence infections; however, the application and impact of quarantine in low prevalence settings appears less common and lacks a formal analysis. Here, we present a quantitative framework using a series of progressively more biologically realistic models of canine rabies in domestic dogs and from dogs to humans, a suitable example system to characterize dynamical changes under varying levels of dog quarantine. We explicitly incorporate health-seeking behaviour data to inform the modelling of contact-tracing and exclusion of rabies suspect and probable dogs that can be identified through bite-histories of patients presenting at anti-rabies clinics. We find that a temporary quarantine of rabies suspect and probable dogs provides a powerful tool to curtail rabies transmission, especially in settings where optimal vaccination coverage is yet to be achieved, providing a critical stopgap to reduce the number of human and animal deaths due to rabid bites. We conclude that whilst comprehensive measures including sensitive surveillance and large-scale vaccination of dogs will be required to achieve disease elimination and sustained freedom given the persistent risk of rabies re-introductions, quarantine offers a low-cost community driven solution to intersectoral health burden.

**Author summary:** Canine rabies remains a human health risk in many countries around the world, particularly in lower and middle income settings where many dogs are free roaming and able to interact more easily with other dogs and humans. In this paper, we present results from a mathematical model that simulates the spread of rabies both between dogs and from dogs to humans and investigate the impact of quarantine and vaccination at reducing transmission. Our work demonstrates the effectiveness of quarantining both infected and exposed dogs - we observe that quarantine can have a substantial effect on reducing the number of new animals subsequently infected and thereby lowering the risk of humans being exposed to infection. Such a policy can have significant benefits, particularly in settings where access to vaccinations is challenging and resources are limited. Our research can therefore help to inform policy makers in countries where canine rabies is circulating to develop appropriate strategies to reduce the human health risks associated with canine rabies in the future.

## Introduction

Canine rabies, an acute zoonotic infection, has been long an enigma in the field of quantitative epidemiology. While deceivingly easy to trace and hence parameterize as transmission happens predominantly among domestic dogs through saliva of an infected individual, model-based predictions are scarcely ever consistent with empirical observations [1]. It is likely due to the complexity and many interdependent factors of the system that the traditional epidemiological models fail to translate to real-world dynamics in their entirety. The details of how rabies transmission operates across temporal and demographic scales has only recently begun to be formally characterized by [2]; however, the broader aspects of the disease epidemiology have been widely explored.

International organizations committed to the global elimination of human deaths from dog-mediated rabies by 2030, and scientific guidance to facilitate progress towards elimination has been underway [3, 4]. Decades of operational experience supported by a mounting body of analytical work conclusively demonstrate that mass vaccination of the dog population is the single most important and cost-effective way to control rabies [5–7]. While implementation of high coverage-achieving, spatially comprehensive annual mass dog vaccination campaigns should be prioritized where possible, the desired control efforts may be impeded by logistical constraints such as availability of resources and limited manpower. Supplementary measures to support vaccination campaigns where coverage (temporarily) falls below the recommended threshold (*<* 70% in [8]) are, however, sparse, and often focused on culling of dogs that has been repeatedly shown ineffective in the case of rabies [6, 9, 10].

While immunization of the susceptible population is the primary intervention strategy in the modern world, quarantine – understood as an isolation of confirmed or probable infected cases – is one of the oldest, low-technology forms of disease control [11, 12]. Transmission potential of an infectious disease is driven by the basic reproduction number (*R*_0_), defined as the average number of secondary cases caused by an infectious individual in a naïve (non immunized) population. *R*_0_ depends on the probability of infection given contact between an infectious and susceptible individual, the length of infectious period, and the number of contacts an infectious individual has per a unit time [13]. Both intervention strategies operate by lowering transmission potential through reducing the number of disease-exposure contacts among hosts.

Vaccination focuses on the reduction of susceptible individuals available to infection and is particularly effective for highly transmissible diseases for which a large proportion of a population would be exposed to the disease agent [14–16]. For infections that circulate endemically at low prevalence, or infections at the early stages of an outbreak, contact-tracing followed by quarantine of probable/infectious individuals provides a highly sensitive tool to curtail the transmission potential of a disease [17, 18]. Classic examples of quarantine measures taken at the onset of an epidemic can be found for outbreaks as old as the bubonic plague pandemic in European port cities and early outbreaks of cholera [19, 20], the 1918 pandemic of influenza [21], to more recent emergencies of Ebola [22], the 2009 A(H1N1)pdm09 influenza [23] and COVID-19 [24]. Conversely, in the case of less-frequent/low-prevalence infections quarantine is commonly applied as a community-based measure especially in low- and middle-income settings, but sparsely implemented as a public health response, possibly due to the lack of formal evidence of its effect.

Here we seek to develop a set of epidemiological models for rabies transmission to examine the effects of quarantine on the disease dynamics. We first focus on the development of an analytical model to explore the qualitative impact of quarantine on the long-term behaviour of the system, specifically interested in analysing the theoretical underpinnings of disease persistence and extinction for low prevalence diseases. Building upon the conceptual understanding gained through the mathematical model, we then expand the existing baseline framework by incorporating probabilistic features relevant to rabies ecology. This allows us to quantitatively investigate the changes in rabies dynamics across varying levels of dog vaccination coverage and under the following quarantine scenarios: (1) no quarantine, (2) quarantine of dogs identified through bite-histories of patients presenting at anti-rabies clinics, and (3) enhanced quarantine informed by contact-tracing of rabies suspect and probable dogs.

## Materials and methods

### Theoretical Model

In order to investigate the long term dynamics of canine rabies within the dog population, we firstly develop a theoretical model that we can utilise to explore stability properties of the system subject to different values of the basic reproduction number (*R*_0_), quarantine rates and vaccination coverage. We consider here an SEIQV model, whereby dogs are either susceptible to infection (*S*), exposed (infected but not yet infectious, *E*), infectious (*I*), quarantined (infectious dogs that are placed in isolation and cannot infect other dogs for the duration of quarantine, *Q*) and vaccinated (*V*).

Note that for rabies we assume that all infected individuals subsequently die from disease. We therefore do not explicitly consider the removed class for this model. The equations governing this system can be defined as follows:

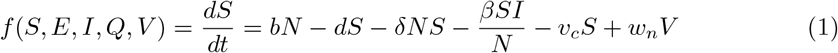

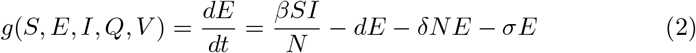

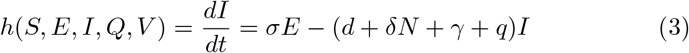

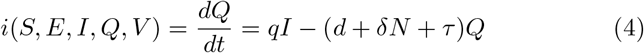

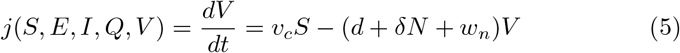

where *N* = *S* + *E* + *I* + *Q* + *V*. In this set of equations, *b* is the birth rate, *d* is the natural death rate, *σ*is the rate of transition from the exposed to the infectious class, *γ* is the death rate from disease, *v*_*c*_ is the vaccination rate, *w*_*n*_ is the rate of waning immunity following vaccination, *τ* is the rate of removal from quarantine and 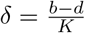 where *K* is the carrying capacity of the population. In contrast to the computational model below in which vaccinated dogs can be re-vaccinated before their immunity wanes, in the deterministic framework only susceptible dogs can be vaccinated.

At epidemic onset and in the absence of quarantine (*q* = 0), we can therefore define the basic reproduction number for this system as

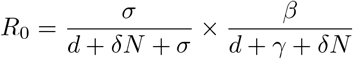

where 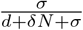 is the fraction of individuals who successfully progress from the exposed to the infectious class and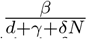 is the transmission rate divided by the average duration that an individual is infectious for.

We will now explore the stability of the system as it approaches equilibrium. Equilibrium solutions occur when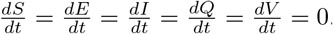 From equations 3 and 4, in equilibrium we find that

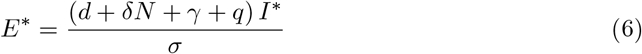

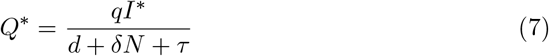

We now substitute our expression for *E*^*∗*^ in equation 6 into equation 2 such that, in the endemic equilibrium (when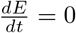and I ?= 0) we find:

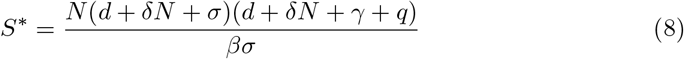

Substituting this expression into equation 5, we can obtain an expression for *V* ^*∗*^ in the endemic equilibrium:

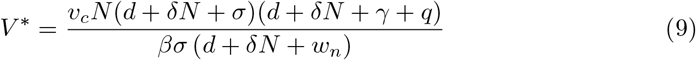

We can now substitute our expressions for *S*^*∗*^ and *V* ^*∗*^ into equation 1 to obtain an expression for *I*^*∗*^:

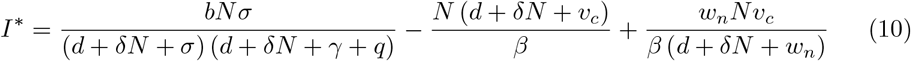

Finally, we can substitute the expression for *I*^*∗*^ into equations 6 and 7 to obtain expressions for *E*^*∗*^ and *Q*^*∗*^:

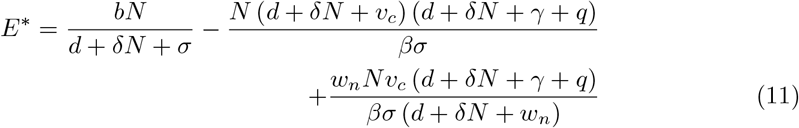

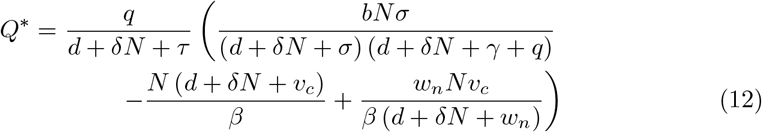

In order to analyse the stability of the system, we need to calculate the Jacobian matrix, *J*. Given that *N* = *S* + *E* + *I* + *Q* + *V* and assuming that *N* is fixed, we can set *V* = *N*− *S*− *E*− *I* −*Q* and reduce the system to consider the four variables *S, E, I* and *Q. J* for this system is therefore

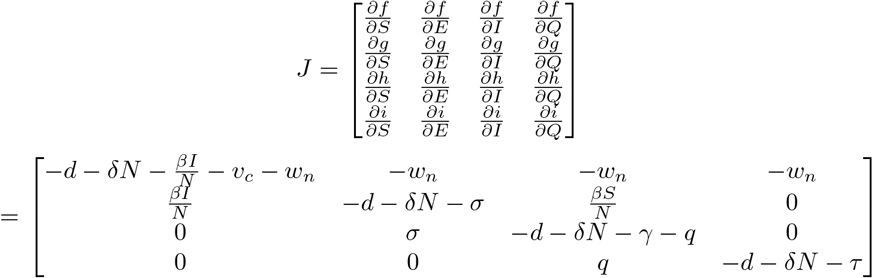

To determine the behaviour of this system, we need to calculate the eigenvalues of the Jacobian. We therefore need to find the solution of |*J* − *λI*| = 0 such that:

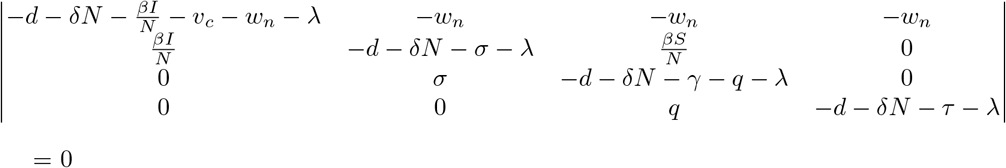

To evaluate the stability of the endemic equilibrium in the context of intervention strategies that might be applied to the system, we use a numerical solving method to calculate the eigenvalues for given values of *R*_0_, vaccination rate *v*_*c*_ and quarantine rate *q*. As such, we determine both when the endemic equilibrium is stable and how the number of infected individuals at the endemic equilibrium depends upon these quantities. A flow diagram of the single-species model is shown in Fig 1. All computational work is performed in the programming environment R, version 3.6.3.

**Fig 1.**
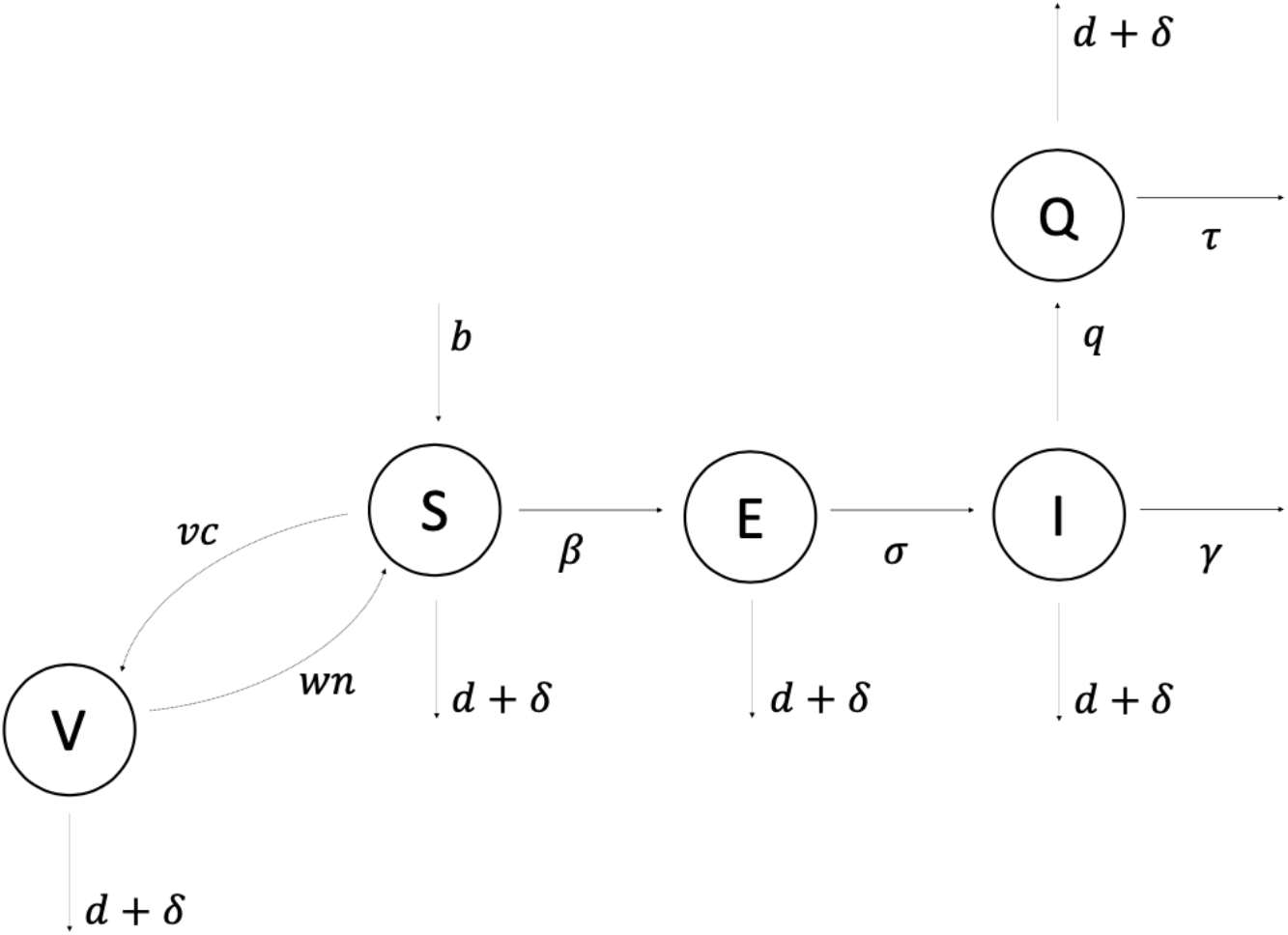
Single-species SEIQV deterministic compartmental model diagram. Epidemiological classes are indicated by circles and arrows suggest the directionality of transitional flows of individuals and the virus moving between compartments. Susceptible individuals can become either Exposed at rate *β* or Vaccinated at rate *v*_*c*_. Vaccinated individuals then return to the Susceptible class with waning immunity of the vaccine at rate *w*_*n*_ which is given by the reciprocal of the average longevity of the vaccine. Infectious individuals can be taken out of their class and placed into Quarantine at rate *q* (note that here we assumed that only Infectious individuals can transition into the Quarantined class). In this scheme, all infectious dogs are removed from the population and die at rate *γ* for Infected individuals and rate *τ* for the Quarantined dogs. Here *τ* = *γ/*(1 −*γρ*) where *ρ* is the mean delay from an individual becoming infectious to entering quarantine, assuming all Quarantined individuals will always die and be removed from the population.

### Computational Model

Whilst the theoretical model presented above provides useful insights regarding the evolution of a rabies-like system in the presence of vaccination and quarantine, transmission of the virus in real-world settings is highly stochastic and can be significantly influenced by low probability events, such as incursions of infected animals into the population or super-spreading events. In fact, stability analyses of deterministic models only consider systems at their fixed points in a closed population, and ignore the role of chance with regards to the pathogen extinction and reintroduction (both locally and globally) despite its influential impact on the future trajectories for diseases that operate at such low transmission levels. For example, the probability of a disease going extinct decreases with an increasing value of *R*_0_ and vice versa [25]. As such, the deterministic threshold for elimination will be modulated by chance processes that can break individual chains of transmission resulting in a faster elimination, or allow the pathogen to persist for longer through a series of infection events and chance re-introductions in spite of an overall high level of immunity within the population.

With this in mind, we now develop a stochastic SEIRQV compartmental model to simulate the spread of disease in the dog population, coupled with an SEIRV model for humans. The system is summarised in Fig 2. In order to introduce a degree of biological realism, we expand the existing SEIQV model by the following additions most relevant from the empirical work. We first (1) introduce the role of stochasticity by modelling both the disease and population dynamics as a probabilistic process, (2) re-define the transmission rate to capture heterogeneity in individual biting behaviour, and (3) allow for exogenous incursions to enter the population. We then introduce an R compartment to keep count of all dogs “Removed” from the population by natural death and the disease (4). We build further realism to modelling the dog quarantine practice, by (5) allowing for a potential removal of Exposed dogs from quarantine when disease symptoms do not occur within the recommended time period of dog exclusion (14 days). Lastly, we extend the single host model to (6) include transmission from dogs to humans, and (7) to incorporate information on health-seeking behaviour and post-exposure prophylaxis (PEP) uptake collected through a longitudinal enhanced surveillance study of dog bite-injury patients [26] in order to approximate the probability of quarantine under different surveillance scenarios.

**Fig 2.**
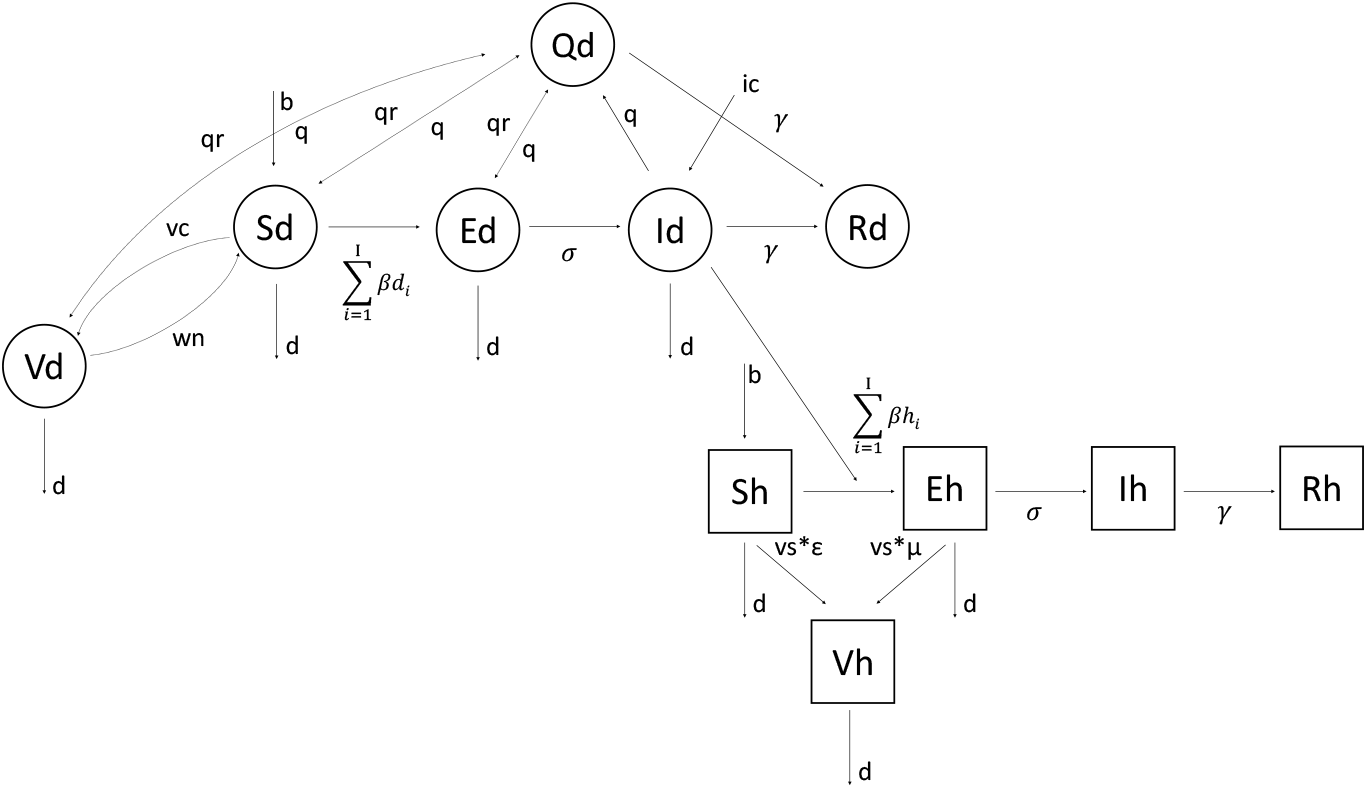
Multi-species SEIRVQ compartmental model diagram. Dog and human epidemiological classes are indicated by circles and squares respectively. Arrows show directions at which individuals and the pathogen move through the system. Compared to the single-species model, here we allow for any epidemiological class of the dog population (except for the Removed class) to be placed in quarantine. Susceptible and Vaccinated quarantined dogs are returned to their respective compartments upon completion of the quarantine at the rate *q*_*r*_. Infected quarantined dogs are removed from the population as a result of disease-induced death at the rate *γ*. Depending on the progression of the disease in Exposed quarantined dogs, two distinct scenarios can occur. For those individuals that will become Infectious within the time frame of their quarantine, disease-induced death follows at the same rate as for Infected individuals, whereas Exposed quarantined dogs that are asymptomatic by the end of their quarantine are returned back into the Exposed class at the rate *q*_*r*_. Transmission of the disease between dogs, and from dogs to humans is defined as a sum of offspring rabid bites seeded by Infected individuals, drawn from a negative binomial distribution taking different parameter values for dogs and human. Lastly, the overall level of infection in the system can be elevated by an introduction of exogenous incursion entering the system at the rate *i*_*c*_. All model parameters associated with the disease and demographic process illustrated here are summarized in Table 1.

Specifically, we model the time stepping process weekly using the Tau leap algorithm.

**Table 1.**
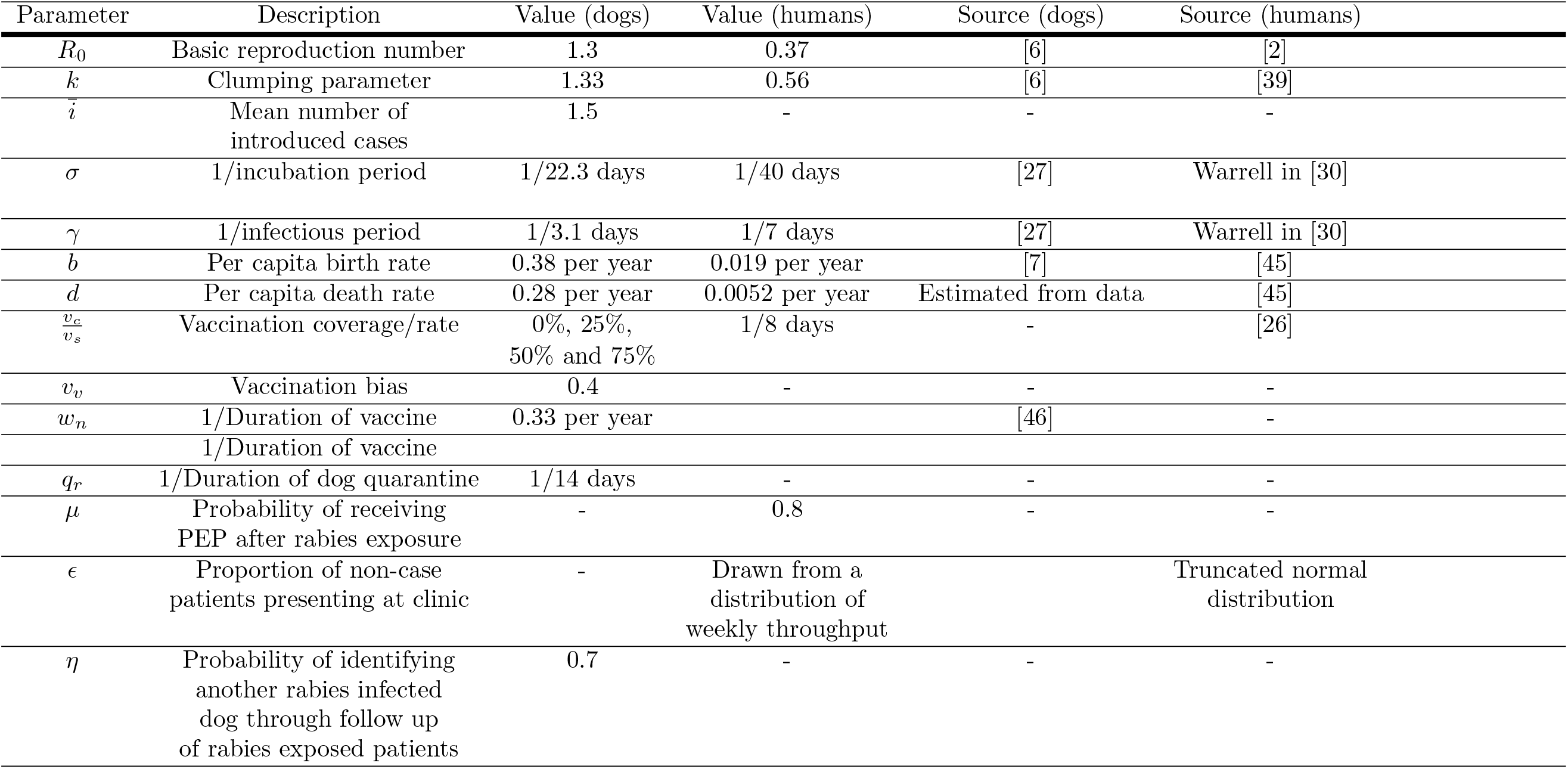
Summary of SEIRVQ model parameters. Parameter values are provided for dogs and humans separately with respective source references.

We then parameterise rabies transmission explicitly as the number of rabid bites per infectious individual. Offspring cases (here representing a secondary case resulting from a biting incident caused by a primary case individual, not a vertical transmission from a parent to its offspring) are drawn from a negative binomial distribution as

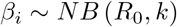

where *R*_0_ is the expected number of new infectious bites (here, 1.3 taken from [2]) and *k* takes different values for humans and dogs. The total transmission rate *β* at each time step is then formulated as a sum of all offspring cases present in the system at the modelled time step, multiplied by the probability of a bite becoming a case (∼50% in [27]), and distributed proportionally to the size of each compartment available to exposure (all except for individuals in Quarantine).

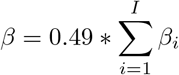

Incursions *i*_*c*_ are drawn from a Poisson distribution where

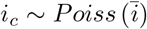

The true rate at which a location receives an incursion will likely vary over time as control is implemented in neighbouring provinces, and geographically given the localized heterogeneous nature of rabies incidence. Here, we incorporate incursions to maintain fluidity in the disease system but set the value to function only as a “background” rate 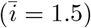

We define dog quarantine as the number of dogs identified through a triage of patients presenting at anti-rabies clinics. The number of quarantined dogs is then drawn from a Conway-Maxwell-Poisson distribution:

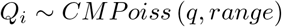

where *q* and *range* differ between investigations of case and non-case incidents. For rabies Exposed and Infected dogs (divided proportionally according to the duration of incubation and infectious periods) identified through patient investigations, the total number of dogs per time step moved into quarantine is then calculated as

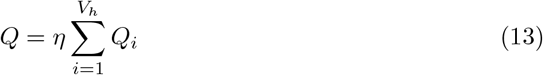

where *η* is the probability that the dogs identified through following rabid animals responsible for exposed case patients are also infected with rabies (i.e., Exposed or Infected). From field observations we believe that *η* is relatively high, but for modelling purposes here we opt for a more conservative assumption of 70% as more data are needed for rigorous estimates. Otherwise, the dogs responsible for non-case incidents both in humans and dogs are distributed in proportion to the size of each relevant compartment (i.e., *S, E, Q*, and *V* within the dog population).

Given the duration of incubation period is longer than the duration of quarantine (22.3 days in [27] and 14 days respectively), a fraction of Exposed quarantined dogs may not become symptomatic before their release. To account for such a possibility, we explicitly generate days until symptomatic for each exposed dog held in quarantine, and return those individuals showing no symptoms after the 14-day period back into the population.

Immunity of humans is defined as achieved through administration of PEP upon attendance at a clinic (note, here we assume that two doses of PEP delivered at days 1 and 7 would provide immunity). The weekly proportion of bite-injury patients *ϵ* is drawn from a zero truncated normal distribution of weekly throughput records collected at the anti-rabies clinics [26]. The percentage of rabies exposed humans that will receive PEP (*µ*) varies extensively across geographical areas and socioeconomic backgrounds. Here, we assume that with enhanced surveillance 80% of human cases would be detected in a timely manner and administered the lifesaving vaccine.

Lastly, we incorporate bias in re-vaccination of dogs (*v*_*v*_) directed towards individuals that are easy to capture for administration of the vaccine. All parameters are summarized in Table 1.

We utilise our computational model to investigate the impact of varying levels of quarantine on rabies dynamics under four vaccination scenarios: 0%, 25%, 50% and 75% of the dog population. We test three progressively strengthened quarantine scenarios. No quarantine is implemented under Scenario 1. In Scenario 2 we assume only dogs identified through bite-injury patients presenting at anti-rabies clinics would be quarantined, suggesting a medium-level quarantine with an average number of dogs per patient (both non-case, and exposed case patients) around 1 (drawn from Conway-Maxwell-Poisson distribution where *q* = 2.5 and *range* = 4.3). Under Scenario 3 we assume a triage of bite-injury patients as per Scenario 2, but this time coupled with further field investigations and contact tracing of patient biting dogs. As such, we expect the number of dogs identified for quarantine through non-case patients to remain within the same range as in Scenario 2, but to increase for investigations informed by exposed case incidents (drawn from Conway-Maxwell-Poisson distribution again where *q* = 2.8 and *range* = 1.5). A computed distribution of the number of dogs identified for quarantine under each of the tested treatments is shown in figure 3. Given the highly stochastic nature of the model, each scenario is iterated 1000 times and run over five consecutive years.

**Fig 3.**
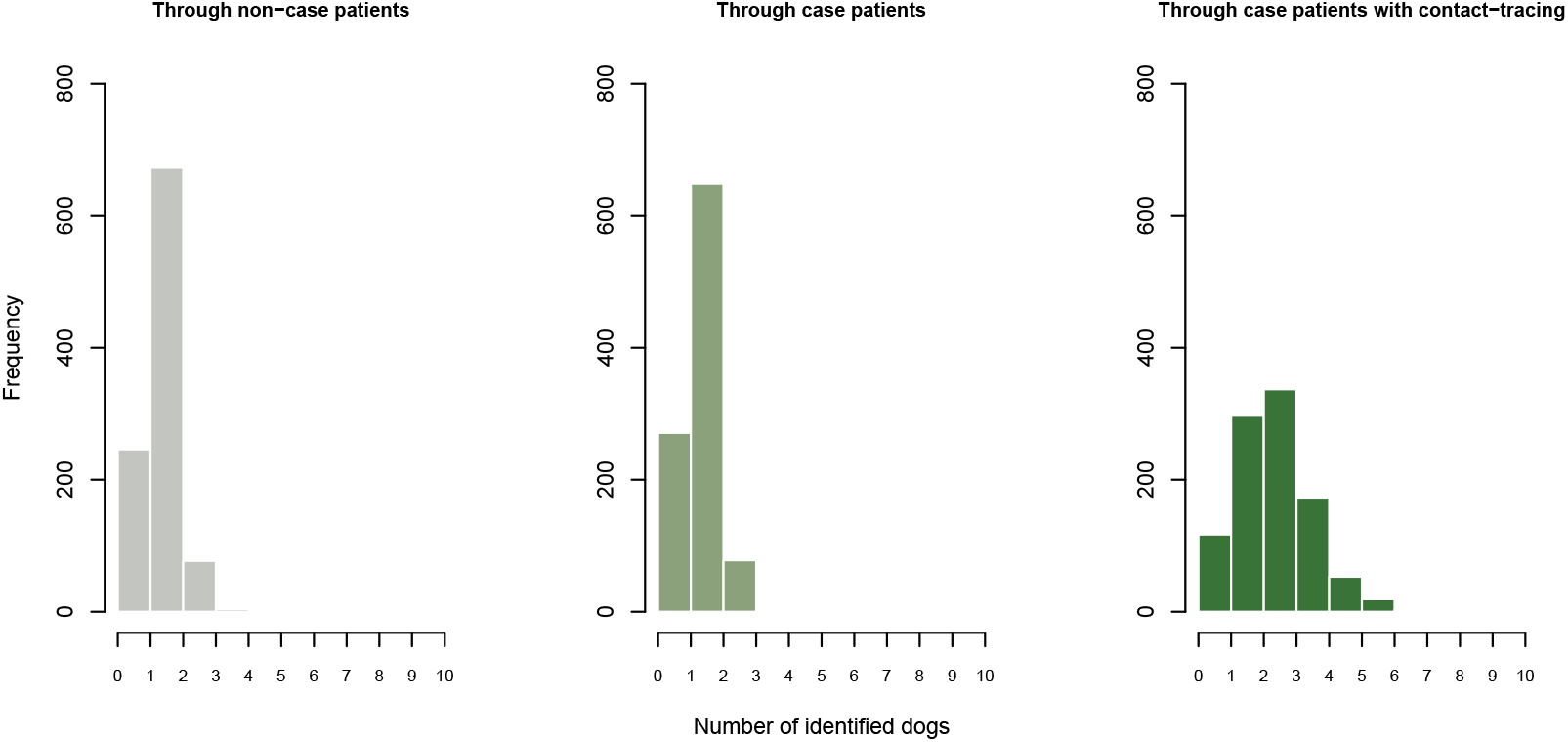
Distributions of the number of dogs identified for quarantine across surveillance scenarios. Frequency distribution of the number of dogs identified for quarantine per biting dog responsible for (from left to right) non-case patients, rabies exposed case patients, and rabies exposed case patients coupled with additional in-field contact tracing investigations of these incidents. Estimates are drawn from a Conway-Maxwell-Poisson distribution with different parameter values taken for each scenario.

## Results

### Stability Analysis of Theoretical Model

Existing models fitted to rabies time-series data suggest that the distribution of *R*_0_ falls predominantly between 1 and 2 [6, 27–29]. As such, we vary the transmission rate *β* by gradually increasing the value of *R*_0_ from 1 to 2 in increments of 0.1 (where *q* = 0 initially to emulate the baseline transmission rate under no intervention). We then set the percentage of infected dogs terminating in quarantine every week (*q*_*p*_) to vary between 0 and 100% in 5% increments, where the rate of quarantine 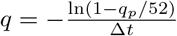 and ∆*t* = 1 (note, all parameters are expressed as weekly rates). To explore the stability of the endemic equilibrium dependent upon given vaccination coverage, quarantine rates and values of *R*_0_, we also vary the mean percentage of dogs vaccinated per year, *v*_*p*_ in 5% increments from 0% to 100%. The weekly vaccination rate is calculated in terms of the percentage of vaccinated dogs such that 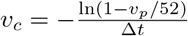where ∆*t* = 1. All remaining parameters used in both models are summarized in Table 1, except for the rate at which Infected dogs leave the quarantine class, which is defined as *τ* = *γ/*(1− *γρ*), where *ρ* is the mean delay from an individual becoming infectious to entering quarantine.

For each combination of parameter values, we then calculate representative eigenvalues to determine the stability of the endemic equilibrium and the size of the infected population (i.e., the number of exposed and infectious dogs) for parameter combinations at which the endemic equilibrium is found to be stable. The results are summarised in figure 4.

**Fig 4.**
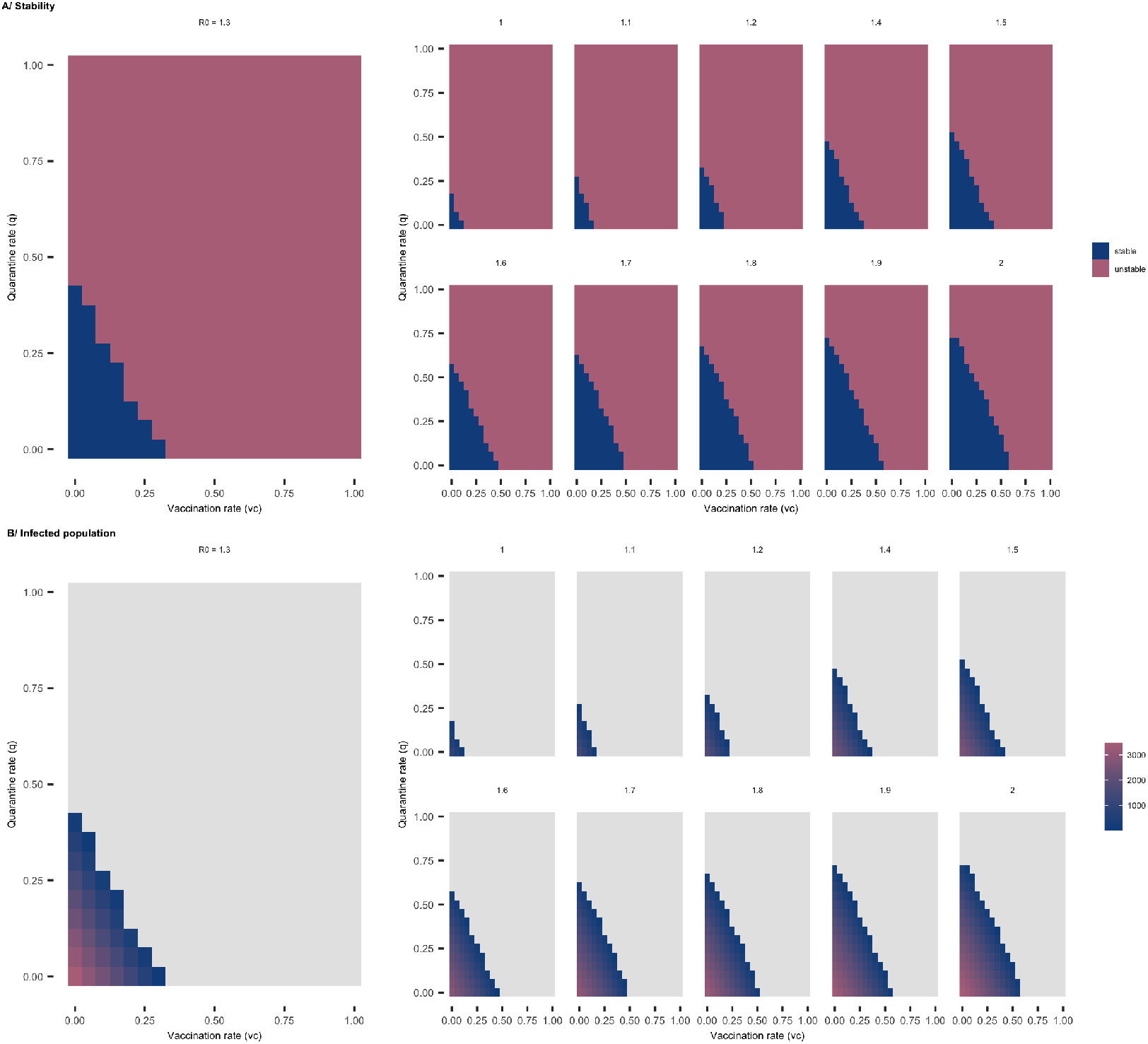
Rabies dynamics across vaccination and quarantine parameter space for different values of *R*_0_. **A**: Stability analysis for the deterministic SEIQV model under *R*_0_ values ranging from 1 to 2. Raster shading shows the stability at the endemic equilibrium for different combinations of vaccination and quarantine rates. The blue shading indicates the region where the endemic equilibrium is stable, whilst the red region indicates where the endemic equilibrium is unstable. **B**: The total number of infected dogs (Exposed and Infected individuals) for vaccination and quarantine rates from 0 to 1 when the system is in endemic equilibrium. In this panel grey shading indicates the region where the disease free equilibrium is stable.

Initially we consider the case of *R*_0_ = 1.3, which previous research indicates as the most likely value for the basic reproduction number for rabies [2]. In the absence of quarantine, we find that as vaccination rates approach 0.35 the endemic equilibrium loses stability and the disease free equilibrium becomes stable. Similarly, in the absence of vaccination, the same result occurs for quarantine rates above 0.45 (figure 4 panel *A*). As the levels of vaccination increase, lower levels of quarantine are required to result in the endemic equilibrium losing stability and the virus being eliminated. As we approach this transition we note that the number of infected dogs in the endemic equilibrium decreases, highlighting the effectiveness of vaccination and quarantine at reducing the number of Infected dogs in the population (figure 4 panel *B*). However, to maintain disease endemicity for *R*_0_ = 1.3 in the absence of both interventions, the required number of Infected dogs (*I*^*∗*^) in the population reaches significantly higher values than what is suggested by empirical evidence (*>* 2110).

We now explore the impact of different values of *R*_0_ upon the stability properties of the endemic equilibrium as vaccination and quarantine rates are varied. When *R*_0_ is close to 1 only very low rates of vaccination and/or quarantine are required in order for elimination to occur. As *R*_0_ increases towards 2, much higher quarantine and vaccination resources are required in order for the endemic equilibrium to become unstable. When *R*_0_ = 2 (which we note represents the upper limit of a realistic value for the basic reproduction number for rabies) we observe that it is possible for rabies to be eliminated provided that sufficient rates of quarantine and vaccination are maintained (figure 4 panel *B*). However, the increased rates of vaccination and particularly quarantine required for elimination in this scenario may be unrealistic and/or infeasible in practice given constrained resources for vaccination, the limited capacity to successfully identify infected dogs for quarantine and the ability to isolate a large number of dogs at any given time.

### Computational Analysis

Any deterministic framing precludes variability in parameter values and the role of chance. In the case of the analytical model, we assume that quarantine, as well as vaccination coverage are maintained consistently over time. We further ignore the duration of exposure which spans a wide temporal range. Symptoms of rabies in dogs usually manifest in the first month since exposure, but the incubation period may last for several months, effectively functioning as an “endogenous” incursion [30, 31]. This becomes particularly relevant when we introduce constantly changing vaccination coverage associated with a build-up of susceptible dogs and a chance of receiving exogenous cases, consequently resulting in variability in transmission.

To address the key limitations of the deterministic framework, we build a stochastic discrete-time multispecies SEIRVQ model with explicit individual biting behaviour and the probability of rabies incursions entering the system from outside. Here we account for the temporal variability in dog vaccination and quarantine, and the impact of chance on the disease dynamics.

In line with the previous results obtained from the deterministic setting, increasing quarantine and vaccination coverage has a positive effect on curtailing the epidemic and leads to significant reductions in the overall number of infections, both in humans and dogs (figure 5 and table 2). However, in the stochastic formulation, neither the vaccination nor the quarantine interventions result in a complete interruption of transmission.

**Table 2.** Trends in the number of dog and human cases under vaccination and quarantine scenarios. All variables statistically significant with *p <<* 0.005. Incident rates for vaccination (continuous) and quarantine (categorical) treatments are calculated from regression coefficients obtained from via negative binomial general linear model.

| $R_0$ | Vaccination<br>Incident<br>Rate (dogs) | Quarantine<br>Scenario 2<br>Incident<br>Rate (dogs) | Quarantine<br>Scenario 3<br>Incident<br>Rate (dogs) | Vaccination<br>Incident<br>Rate (humans) | Quarantine<br>Scenario 2<br>Incident<br>Rate (humans) | Quarantine<br>Scenario 3<br>Incident<br>Rate (humans) |
| --- | --- | --- | --- | --- | --- | --- |
| 1.0 | 0.302 | 0.834 | 0.643 | 0.605 | 0.875 | 0.797 |
| 1.1 | 0.268 | 0.825 | 0.628 | 0.563 | 0.858 | 0.772 |
| 1.2 | 0.235 | 0.815 | 0.613 | 0.510 | 0.846 | 0.753 |
| 1.3 | 0.204 | 0.798 | 0.593 | 0.462 | 0.832 | 0.729 |
| 1.4 | 0.174 | 0.779 | 0.571 | 0.409 | 0.813 | 0.704 |
| 1.5 | 0.147 | 0.756 | 0.543 | 0.351 | 0.785 | 0.670 |
| 1.6 | 0.120 | 0.732 | 0.521 | 0.299 | 0.752 | 0.638 |
| 1.7 | 0.093 | 0.687 | 0.480 | 0.232 | 0.699 | 0.577 |
| 1.8 | 0.069 | 0.628 | 0.431 | 0.171 | 0.633 | 0.505 |
| 1.9 | 0.045 | 0.535 | 0.359 | 0.103 | 0.519 | 0.404 |
| 2.0 | 0.023 | 0.380 | 0.251 | 0.046 | 0.358 | 0.265 |

**Fig 5.**
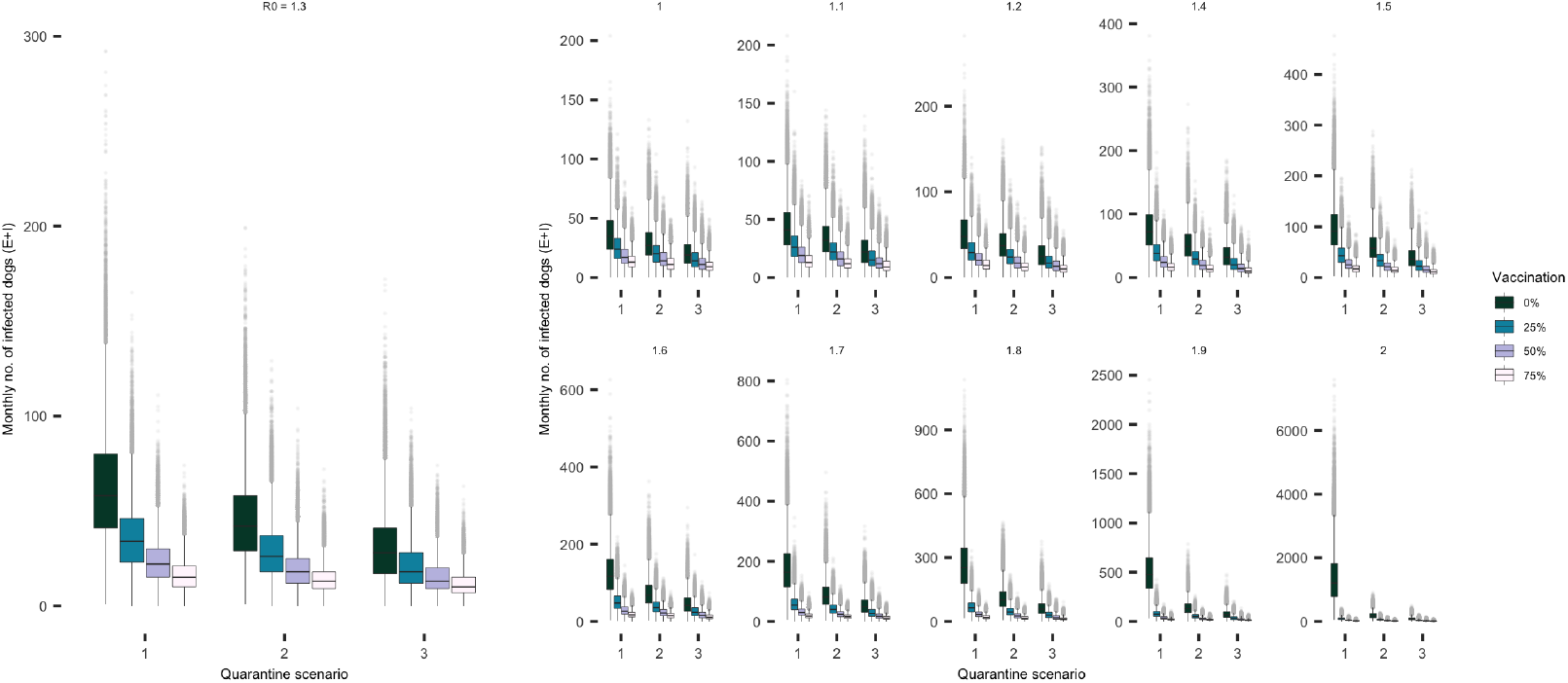
Incremental decline in the monthly number of total infection (i.e., Exposed and Infected dogs) in the dog population across all 1000 simulations with increasing levels of dog quarantine and vaccination coverage. Summaries shown as individual panels for a range *R*_0_ values (please note the differential scale of y-axes between the panels). Differences between quarantine scenarios in the number of Exposed and Infected dogs is particularly striking in low vaccination coverage settings and/or for higher values of *R*_0_ as supported by the statistical analysis summarized in Table 2. Note, the simulation data points do not include the initial burn-in period of the first six months.

To probe the interaction between vaccination and quarantine measures, we test four incrementally increasing levels of vaccination coverage (i.e., 0%, 25%, 50% and 75%. Whilst for the deterministic framework, *>* 20% vaccination coverage was found to be sufficient in order to drive the system to extinction in the presence of low-level quarantine (figure 4 panel *A*), this threshold is likely inaccurate for a system in which the vaccination coverage changes over time as a result of variable vaccination rate, fast turnover of susceptible individuals through high birth and death rates, and variability in the number of offspring cases for each infectious dog. The effects of quarantine (both medium and high level - scenarios 2 and 3 respectively) on the number of infections in the population appear particularly remarkable in no- and low-vaccination settings, with lesser impact on the system as the vaccination coverage increases (figure 5 and 6).

**Fig 6.**
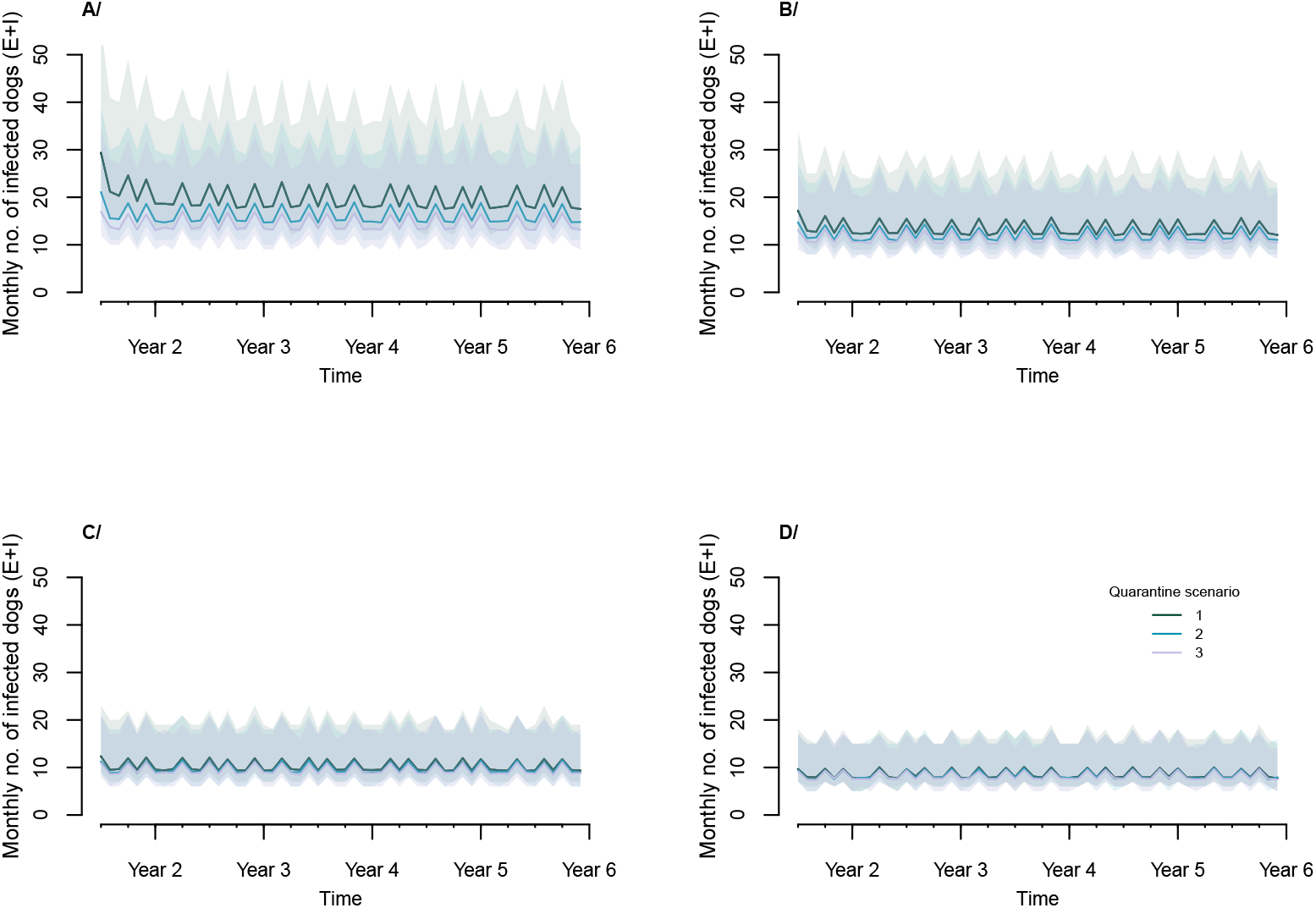
Example time series for *R*_0_ = 1.3, demonstrating rabies dynamics under quarantine scenarios across vaccination coverage levels. Rabies levels circulating in dogs when no quarantine is implemented shown in dark green (Scenario 1), for medium-level quarantine (Scenario 2) in blue, and under intensified quarantine informed by contact-tracing (Scenario 3) in purple. Shading around projected trajectories indicate the 95% confidence envelope. Note that the time series omit the model burn-in period of the initial six months. **A**/no vaccination coverage. **B**/25% vaccination coverage. **C**/50% vaccination coverage. **D**/75% vaccination coverage.

However, introductions of rabies cases from outside the population pose an additional impediment to disease elimination. Exogenous incursions increase the magnitude of transmission temporarily and decrease the probability of extinction in the long term. In fact, under increased detection and quarantine of infected dogs through field investigations (Scenario 3) and the intensified vaccination efforts, endemicity appears to be sustained predominantly through incursions. Similar dynamics have been reported for diseases with lower transmission rates [32] and/or during the endgame (i.e. pre-elimination/pre-eradication epidemiological stage as described in [33, 34], when heterogeneities in the force of infection (often driven by incursions) and the level of immunity may result in unpredictable stochastic outbreaks [33, 35].

## Discussion

Canine rabies circulating in domestic dogs represents a serious burden on public health budgets and local communities. Mass dog vaccination campaigns, the cornerstone of effective rabies control, has led to elimination of human deaths and interruption of rabies transmission at the source in most countries across the Global North [36]. Such campaigns, however, require systematic efforts delivered at scale and sustained over long periods of time [6]. As the availability of human and financial resources is typically limited in low- and middle-income countries, questions remain over the most effective strategies to eliminate rabies given extensive technical and structural constraints.

Contact tracing and subsequent quarantine of infectious individuals plays an important role in the control of infectious diseases at the onset of an outbreak or during the endgame [37, 38]. Concentrating control on infected contacts can be potentially extremely effective, but it relies on a sensitive surveillance system and a logistically traceable fraction of the infected population. Such conditions are typically met during the early or final stages of an epidemic when only a limited number of infections is present within the system, and for diseases with easily recognizable symptoms.

Whilst endemic, rabies provides a unique system to test a wider use of quarantine outside its traditional application. Biting behaviour is the primary indicator of rabies; transmission events, particularly from dogs to humans, are extremely memorable as often inducing severe distress or even psychological trauma. Thus, they are relatively easy to identify (when investigated) and traced back and forward as the local communities remember the bite histories long after they have occurred. In addition, rabies circulates at a low prevalence with *R*_0_ *<* 2, indicating only a small percentage of the population is being infected at any time [27, 39].

The concept of quarantine for rabies suspect and probable dogs has long been part of the general guidelines for community-based rabies control, particularly in low- and middle-income counties. The dynamical impact of such an intervention has, however, never been formally assessed. Using a combination of mathematical and computational models developed to capture rabies dynamics in the context of control interventions guided by health-seeking behaviour, here we investigate the impact of quarantine and vaccination on the stability of the system with potential application to other low incidence diseases.

We found that in the deterministic settings even medium levels of quarantine of infected dogs pose a strong pressure on the stability of the system, and in combination with minimal vaccination efforts quarantine would lead to a complete elimination of the disease. Analytical models are powerful tools to explore global dynamics of a system and its long-term evolution, but the insights are relevant only when assessed qualitatively. As such, the theoretical findings suggest that introducing formal quarantine into the rabies management strategies alongside mass dog vaccination campaigns would result in a reduction in the overall burden of rabies cases whilst potentially providing a critical stopgap in areas where immunity coverage falls temporarily below the optimal levels. However, the exact parameter thresholds for when elimination can be expected are only conceptual and will be modulated in empirical settings.

In fact, in the expanded probabilistic SEIRVQ framework the temporal exclusion of dogs through quarantine does not result in elimination of the pathogen in spite of higher vaccination coverage levels than suggested by the analytical model. Stochastic extinctions of individual transmission chains offset by re-introductions of rabies from outside the population via exogenous incursions create a highly non-linear landscape of transmission, requiring more extensive efforts than predicted deterministically given the highly probabilistic nature of individual transmission events (e.g. barriers to achieving elimination of polio in [33]). Demography may also play an important role; fast population turnover due to high vital rates leads to constant restructuring of the dog population and its immunity profile. Indeed, in areas where the dog population undergoes a substantial demographic change, annual high coverage achieving (*>* 70%) mass dog vaccination campaigns are essential to countervail the immunity loss due to removal of vaccinated dogs and their replacement with susceptible puppies [5, 6].

It is, however, important to note, that the depletion of the susceptible population is not associated with rabies, suggesting that changes in the size of the dog population alone will not affect rabies transmission unless accompanied by additional measures in a holistic manner. Both empirical data and models indicate that rabies transmission between dogs occurs independently of the population density under a vast range conditions, meaning that regardless of the dog population size rabid dogs will produce on average the same number of infectious contacts [2, 6, 9].

Conversely, contacts leading to disease transmission are largely context specific, and they will change as the interventions are being implemented and in response to the phase of the epidemic curve. Social, cultural, environmental and incidental backgrounds can vary widely even across small spatial ranges, resulting in many loosely connected metapopulations that act, for most time, as individual foci [40, 41]. For diseases with higher transmission rates, smaller scale differences can be averaged across larger spatial aggregates/population, whilst for the lower incidence infections detailed spatial models provide partial leverage in capturing some of the system’s heterogeneities. Extensive spatial models can, however, end up being extremely costly and intractable in terms of deriving generalizable results across settings. For example, a variability in the incubation and infectious period distributions can dramatically alter the characteristics of rabies outbreaks which in turn will largely change parameter estimates for each model or setting [40]. Nevertheless, such insights into broader mechanisms of rabies transmission will likely prompt further investigations into the biological drivers of variation in individual biting behaviour beyond population-level factors, that is yet to be captured formally in a mathematical framework.

Our findings add onto the existing body of information on rabies management including actionable guidelines and tools supported by decades of operational research, and offer a deeper understanding of the principles and effectiveness of quarantine on rabies dynamics. Implementing contact tracing and quarantine of suspect and probable dogs may bring enormous benefits to public health and the affected communities, particularly in lower vaccination settings, directly reducing the number of deaths due to rabid bites. However, while active investigations and quarantine appear a powerful component of the One Health response in curtailing transmission, large-scale vaccination of dogs is necessary for complete interruption of transmission of the virus and sustained elimination of rabies, given the enduring risk of re-introductions from neighbouring populations [42–44]. With the aspiration to eliminate dog-mediated human rabies by 2030, we conclude that a successful outcome depends on a combination of complementary intersectoral control measures, integrating and building upon operational capacities of both public health and veterinary sectors.

## Acknowledgments

This work was supported by the Biotechnology and Biological Sciences Research Council the Engineering and Physical Sciences Research Council (BBSRC BB/M01116X/1; EPSRC GCRF EP/R512916/1). We are especially grateful to Ed Hill for useful feedback and comments on the manuscript and figures. Last but not least, we would like to thank Brad, SB, Clint, Clay, AJ and the rest of the Cordova group for providing a solid support network when needed.

## References

1. Rajeev M, Metcalf CJE, Hampson K. Modeling canine rabies virus transmission dynamics. Fooks, A and Jackson, A ed Rabies (Fourth Edition), Academic Press. 2020; p. 655–670.

2. Mancy R, Rajeev M, Lugelo A, Brunker K, Cleaveland S, Ferguson EA, et al. Rabies shows how scale of transmission can enable acute infections to persist at low prevalence. Science. 2022;376:512–516.

3. Minghui R, Stone M, Semedo M, Nel L. New global strategic plan to eliminate dog-mediated rabies by 2030. The Lancet Global Health. 2018;6(8):e828–e829.

4. WHO. Zero by 30: the global strategic plan to end human deaths from dog-mediated rabies by 2030.; 2018. Available from: https://www.who.int/rabies/resources/9789241513838/en/.

5. Coleman P, Dye C. Immunization coverage required to prevent outbreaks of dog rabies. Vaccine. 1996;14(3):185–186.

6. Townsend SE, Sumantra IP, Pudjiatmoko, Bagus GN, Brum E, Cleaveland S, et al. Designing Programs for Eliminating Canine Rabies from Islands: Bali, Indonesia as a Case Study. PLOS Neglected Tropical Diseases. 2013;7(8):e2372.

7. Ferguson E, Hampson K, Cleaveland S, Consunji R, Deray R, Friar J, et al. Heterogeneity in the spread and control of infectious disease: consequences for the elimination of canine rabies. Scientific Reports. 2015;5(1):18232.

8. WHO. WHO Expert Consultation on Rabies. WHO technical report series: Second Report;982.

9. Morters M, Restif O, Hampson K, Cleaveland S, Wood J, Conlan A. Evidence based control of canine rabies: a critical review of population density reduction. Journal of Animal Ecology. 2012;82(1):6–14.

10. Purwo Suseno P, Rysava K, Brum E, De Balogh K, Ketut Diarmita I, Fakhri Husein W, et al. Lessons for rabies control and elimination programmes: a decade of One Health experience from Bali, Indonesia. Revue Scientifique et Technique de l’OIE. 2019;38(1):213–224.

11. Keeling M, Rohani P. Modeling Infectious Diseases in Humans and Animals. Princeton University Press; 2008.

12. Conti AA. Quarantine through history. International Encyclopedia of Public Health. 2008; p. 454.

13. Bjørnstad O. Epidemics: Models and Data Using R; 2018.

14. Bansal S, Pourbohloul B, Meyers LA. A Comparative Analysis of Influenza Vaccination Programs. PLOS Medicine. 2006;3(10):e387.

15. Glasser J, Feng Z, Omer S, Smith P, Rodewald L. The effect of heterogeneity in uptake of the measles, mumps, and rubella vaccine on the potential for outbreaks of measles: a modelling study. The Lancet Infectious Diseases. 2016;16(5):599–605.

16. Lee S, Chowell G. Exploring optimal control strategies in seasonally varying flu-like epidemics. Journal of Theoretical Biology. 2017;412:36–47.

17. Klinkenberg D, Fraser C, Heesterbeek H. The Effectiveness of Contact Tracing in Emerging Epidemics. PLOS One. 2006;1(1):e12.

18. Berge T, Ouemba Tassé A, Tenkam H, Lubuma J. Mathematical modeling of contact tracing as a control strategy of Ebola virus disease. International Journal of Biomathematics. 2018;11(07):1850093.

19. Morris RJ. Cholera 1832: The social response to an epidemic. vol. 20. Routledge; 2022.

20. Beckmann J. A history of inventions, discoveries, and origins. Musaicum Books; 2021.

21. Peltier M. The influenza epidemic that occurred in New Caledonia in 1921. Bull de l’Office Int d’Hygiene Publique. 1922;6:677–685.

22. Pandey A, Atkins KE, Medlock J, Wenzel N, Townsend JP, Childs JE, et al. Strategies for containing Ebola in West Africa. Science. 2015;346(6212):991–995.

23. Schoch-Spana M, Bouri N, Rambhia KJ, Norwood A. Stigma, health disparities, and the 2009 H1N1 influenza pandemic: how to protect Latino farmworkers in future health emergencies. Biosecurity and bioterrorism: biodefense strategy, practice, and science. 2010;8(3):243–254.

24. for Disease Control C, Rothstein MA, Alcalde MG, Elster NR, Majumder MA, Palmer LI, et al. Quarantine and isolation: Lessons learned from SARS. University of Louisville School of Medicine, Institute for Bioethics, Health … ; 2003.

25. Lloyd-Smith J, Schreiber S, Kopp P, Getz W. Superspreading and the effect of individual variation on disease emergence. Nature. 2005;438(7066):355–359.

26. Rysava K, Espineda J, Silo EAV, Carino S, Aringo AM, Bernales RP, et al. One Health Surveillance for Rabies: A Case Study of Integrated Bite Case Management in Albay Province, Philippines. Frontiers in Tropical Diseases. 2022;3. doi:10.3389/fitd.2022.787524.

27. Hampson K, Dushoff J, Cleaveland S, Haydon D, Kaare M, Packer C, et al. Transmission Dynamics and Prospects for the Elimination of Canine Rabies. PLOS Biology. 2009;7(3):e1000053.

28. Kurosawa A, Tojinbara K, Kadowaki H, Hampson K, Yamada A, Makita K. The rise and fall of rabies in Japan: A quantitative history of rabies epidemics in Osaka Prefecture, 1914–1933. PLOS Neglected Tropical Diseases. 2017;11(3):e0005435.

29. Cori A, Nouvellet P, Garske T, Bourhy H, Nakouné E, Jombart T. A graph-based evidence synthesis approach to detecting outbreak clusters: An application to dog rabies. PLOS Computational Biology. 2018;14(12):e1006554.

30. Kaplan C, Turner G, Warrell D, Brown F, Crick J, Haig D, et al. Rabies THE FACTS. Oxford University Press; 1977.

31. Hemachudha T, Laothamatas J, Rupprecht CE. Human rabies: a disease of complex neuropathogenetic mechanisms and diagnostic challenges. The Lancet Neurology. 2002;1(2):101–109.

32. Bouri N, Sell TK, Franco C, Adalja AA, Henderson DA, Hynes NA. Return of epidemic dengue in the United States: implications for the public health practitioner. Public Health Reports. 2012;127(3):259–266.

33. Klepac P, Metcalf C, McLean A, Hampson K. Towards the endgame and beyond: complexities and challenges for the elimination of infectious diseases. Philosophical Transactions of the Royal Society B: Biological Sciences. 2013;368(1623):20120137.

34. Barret S. Economic considerations for the eradication endgame. Philosophical Transactions of the Royal Society London B: Biological Sciences. 2013;368(1623):20120149. doi:10.1098/rstb.2012.0149.

35. Grassly N, Fraser C. Mathematical models of infectious disease transmission. Nature Reviews Microbiology. 2008;6(6):477–487.

36. Taylor L, Nel L. Global epidemiology of canine rabies: past, present, and future prospects. Veterinary Medicine: Research and Reports. 2015; p. 361.

37. Fenner F, Henderson DA, Arita I, Jezek Z, Ladnyi ID. Smallpox and its eradication. WHO History of international public health. 1988;6.

38. Donnelly C, Ghani A, Leung G, Hedley A, Fraser C, Riley S, et al. Epidemiological determinants of spread of causal agent of severe acute respiratory syndrome in Hong Kong. The Lancet. 2003; p. 1761–1766.

39. Hampson K, Abela-Ridder B, Brunker K, Bucheli STM, Carvalho M, Caldas E, et al. Surveillance to Establish Elimination of Transmission and Freedom from Dog-mediated Rabies. BioRxiv. 2016;doi:https://doi.org/10.1101/096883.

40. Beyer H, Hampson K, Lembo T, Cleaveland S, Kaare M, Haydon D. Metapopulation dynamics of rabies and the efficacy of vaccination. Proceedings of the Royal Society B: Biological Sciences. 2010;278(1715):2182–2190.

41. Beyer H, Hampson K, Lembo T, Cleaveland S, Kaare M, Haydon D. The implications of metapopulation dynamics on the design of vaccination campaigns. Vaccine. 2012;30(6):1014–1022.

42. Windiyaningsih C, Wilde H, Meslin FX, Suroso T, Widarso HS. The rabies epidemic on Flores Island, Indonesia (1998-2003). Journal of Medical Association of Thailand. 2004;87:1389–1393.

43. Putra AAG, Hampson K, Girardi J, Hiby E, Knobel D, Mardiana IW, et al. Response to a rabies epidemic, Bali, Indonesia, 2008–2011. Emerging Infectious Diseases. 2013;19(4):648–651.

44. Bamaiyi PH. 2015 Outbreak of Canine Rabies in Malaysia: Review, Analysis and Perspectives. Journal of Veterinary Advances. 2015;5(12):1181.

45. Philippine Statistics Authority: Highlights of the Philippine Population 2015 Census of Population;. http://psa.gov.ph/content/highlights-philippine-population-2015-census-population.

46. Lakshmanan N, Gore TC, Duncan KL, Coyne MJ, Lum MA, Sterner FJ. Three-year rabies duration of immunity in dogs following vaccination with a core combination vaccine against canine distemper virus, canine adenovirus type-1, canine parvovirus, and rabies virus. Veterinary Therapeutics. 2006;7(3):223–231.

